# Desensitization of NMDA receptors depends of their association with plasma membrane sodium-calcium exchangers in lipid nanoclasters

**DOI:** 10.1101/378026

**Authors:** Dmitry A. Sibarov, Ekaterina E. Poguzhelskaya, Sergei M. Antonov

## Abstract

The plasma membrane Na^+^/Ca^2+^-exchanger (NCX) has recently been shown to regulate Ca^2+^-dependent *N*-methyl-d-aspartate receptor (NMDAR) desensitization, suggesting tight interaction of NCXs and NMDARs in lipid nanoclaster or “rafts”. To evaluate possible role of this interaction we studied effects of Li^+^ on NMDA-elicited whole-cell currents and Ca^2+^ responses of rat cortical neurons *in vitro* before and after cholesterol extraction by methyl-β-cyclodextrin (MβCD). Substitution Li^+^ for Na^+^ in the external solution caused a concentration-dependent decrease of steady-state NMDAR currents from 440 ± 71 pA to 111 ± 29 pA in 140 mM Na^+^ and 140 mM Li^+^, respectively. Li^+^ inhibition of NMDAR currents disappeared in the absence of Ca^2+^ in the external solution (Ca^2+^-free), suggesting that Li^+^ enhanced Ca^2+^-dependent NMDAR desensitization. Whereas the cholesterol extraction with MβCD induced NMDAR current decrease to 136 ± 32 pA in 140 mM Na^+^ and 46 ± 15 pA in 140 mM Li^+^, the IC_50_ values for the Li^+^ inhibition were similar (about 44 mM Li^+^) before and after this procedure. In Ca^2+^-free Na^+^ solution steady-state NMDAR currents after the cholesterol extraction were 47 ± 6 *%* of control currents. Apparently this amplitude decrease was not Ca^2+^-dependent. In 1 mM Ca^2+^ Na^+^ solution the Ca^2+^-dependent NMDAR desensitization was greater when cholesterol was extracted. Obviously, this procedure promoted its development. In agreement, Li^+^ and KB-R7943, an inhibitor of NCX, both considerably reduced NMDAR-mediated Ca^2+^ responses. The cholesterol extraction itself caused a decrease of NMDAR-mediated Ca^2+^ responses and, in addition, abolished the effects of Li^+^ and KB-R7943. Taken together our data suggest that NCXs downregulate the Ca^2+^-dependent NMDAR desensitization. Most likely, this is determined by co-localization and tight functional interaction of NCX and NMDAR molecules in membrane lipid rafts. Their destruction is accompanied by an enhancement of NMDAR desensitization and a loss of NCX-selective agent effects on NMDARs.

## Introduction

*N*-methyl-d-aspartate activated glutamate receptors (NMDARs) are ligand gated ion channels which naturally transfer currents determined by Na^+^, K^+^ and Ca^2+^ permeation. High permeability of NMDARs to Ca^2+^ makes them involved in synaptic plasticity [1, 2], while their hyperactivation during ischemia or stroke causes neuronal Ca^2+^ overload and apoptosis [3]. Ca^2+^-dependent desensitization of NMDARs represents a feedback regulation of the NMDAR open probability by the Ca^2+^ entry into neurons [4-8]. The Ca^2+^ entry via NMDAR pores produces a local increase of Ca^2+^ concentration (up to micromolar range) in a close proximity of receptor intracellular domains. Calmodulin binds free Ca^2+^ and then interacts with C-terminal domains of NMDAR GluN1 subunits causing the decrease of the channel open probability in the Ca^2+^ concentration-dependent manner because of Ca^2+^-dependent NMDAR desensitization [9].

Recently it has been demonstrated that the inhibition of the plasma membrane Na^+^/Ca^2+^ exchanger (NCX) either by KB-R7943 (2-[2-[4-(4-nitrobenzyloxy) phenyl]ethyl] isothiourea methanesulfonate) or by the substitution of Li^+^ for Na^+^ in the external physiological solution considerably enhances the Ca^2+^-dependent desensitization of NMDARs [10]. As Li^+^ is a substrate inhibitor of Na^+^-dependent neurotransmitter transporters [11, 12] and exchangers [for review see 13] the substitution of Li^+^ for Na^+^ in the external solution decreases the efficacy of Ca^2+^ extrusion via NCX. The direct effects of Li^+^ on NMDAR kinetics and conductance is negligible, because NMDARs have similar Li^+^ and Na^+^ channel permeabilities [14]. These observations suggest that NCX is involved in regulation of Ca^2+^-dependent desensitization of NMDARs that could be achieved in the case of close location and interaction of NCX and NMDAR molecules in the plasma membrane.

The Li^+^ therapy is widely used to stabilize mood disorders, including bipolar disorders and depression as well as suicidal behaviors [12]. There are some experimental indications that KB-R7943 reduces 4-aminopyridine-induced epileptiform activity in adult rats [15]. It is still not clear whether NCXs could represent a target of pharmacological action to compensate NMDAR-related neuronal pathologies and whether an acceleration of Ca^2+^-dependent NMDAR desensitization by Li^+^ is at least partially contributed to the Li^+^ therapeutic effects. To provide more clues for understanding of these aspects of the NMDAR pharmacology here we study the concentration dependence of Li^+^ effects on NMDAR currents and the role of functional interaction between NCXs and NMDARs that presumably requires their close spatial localization in lipid rafts.

## Materials and methods

### Primary Culture of Cortical Neurons

The procedure of culture preparation from rat embryos was previously described [16, 17]. All procedures using animals were in accordance with recommendations of the Federation for Laboratory Animal Science Associations and approved by the local Institutional Animal Care and Use Committees. Wistar rats (provided by the Sechenov Institute’s Animal Facility) 16 days pregnant (overall 12 animals in this study) were sacrificed by CO2 inhalation. Fetuses were removed and their cerebral cortices were isolated, enzymatically dissociated and used to prepare primary neuronal cultures. Cells were used for experiments after 10–15 days in culture [17, 18]. Cells were grown in Neurobasal^™^ culture media supplemented with B-27 (Gibco-Invitrogen, UK) on glass coverslips coated with poly-d-lysine.

### Patch Clamp Recordings

Whole-cell currents were recorded on rat cortical neurons in primary culture (10–15 days in *vitro)* by patch clamp technique using a MultiClamp 700B amplifier with Digidata 1440A acquisition system controlled by pClamp v10.2 software (Molecular Devices, Sunnyvale, CA, USA) at room temperature (23– 25°C). The signal was 8-order low-pass Butterworth filtered at 200 Hz to remove high frequency noise. Acquisition rate was 20000 s^-1^. Micropipette positioning was performed with an MP-85 micromanipulator (Sutter Instrument, Novato, CA, USA) under visual control using a Nikon Diaphot TMD microscope (Nikon, Japan). For fast medium exchange we used a BPS-4 fast perfusion system (ALA Scientific Instruments, Farmingdale,

NY, USA). The tip of the multichannel manifold was placed at a distance of 0.2 mm from the patched cell, allowing solution exchange in 80 ms. Unless otherwise specified, the following extracellular medium was used for recording (external bathing solution, in mM): 140 NaCl; 2.8 KCl; 1.0 CaCl_2_; 10 HEPES, at pH 7.2–7.4. The patch-pipette solution contained (in mM): 120 CsF, 10 CsCl, 10 EGTA, and 10 HEPES. In some experiments BAPTA ((1,2-bis(*o*-aminophenoxy)ethane-*N,N,N’N’*-tetraacetic acid) was added to path-pipette solution to prevent calcium-dependent desensitization of NMDARs. This solution contained (in mM): 120 CsF, 10 CsCl, 10 EGTA, 10 HEPES, 0.1 CaCl_2_, 1 BAPTA to achieve calculated free [Ca^2+^] of 13 nM. The pH was adjusted to 7.4 with CsOH. Measured osmolarities of the external bathing solution and the patch-pipette solution were 310 and 300 mOsm, respectively. Patch pipettes (2–4 MΩ) were pulled from 1.5-mm (outer diameter) borosilicate standard wall capillaries with inner filament (Sutter Instrument, Novato, CA, USA). In whole-cell configuration the series resistances did not exceed 10 MΩ. Holding membrane voltage (V_m_) was corrected for the liquid junction potential between the Na^+^-containing external bathing solution and the Cs^+^-containing pipette solution of -15 mV.

### Loading of Fluo-3 AM and Ca^2+^ Imaging

Cells were loaded with Fluo-3 AM (4 mM, Life Technologies, Foster City, CA, USA) using conventional protocols as recommended by the manufacturer. In brief, neuronal cultures were incubated with Fluo-3 AM for 45 min in the dark at room temperature. Then, Fluo-3 AM was washed out, and cells were incubated in the external solution for another 30 min in the dark. Coverslips with Fluo-3-loaded neurons were placed in the perfusion chamber, which was mounted on the stage of a Leica TCS SP5 MP inverted microscope (Leica Microsystems, Germany). Fluorescence was activated with 488 nm laser light and emission was measured within the wavelength range from 500 to 560 nm. Images were captured every 1.5 second during 30 min experiments.

### Drugs

Functional activity of NMDARs requires binding of both glutamate and a co-agonist, glycine. Unless otherwise stated, to activate NMDARs we applied 100 μM NMDA with 10 μM L-glycine (Gly). KB-R7943 (2-[4-[(4-nitrophenyl)methoxy]phenyl]ethyl ester, methanesulfonate, 10 μM) application or proportional substitution of Li^+^ for Na^+^ in the external bathing solution were used to inhibit NCX. Methyl-β-cyclodextrin (MβCD, 1.5 mM) application for 5 minutes was used to destruct lipid rafts by extracting cholesterol from the plasma membrane. All compounds were from Sigma-Aldrich, St. Louis, MO, USA or Tocris Bioscience, UK.

### Data Analysis

Quantitative data are expressed as mean ± SEM. ANOVA and Bonferroni multiple comparison methods as well as Student’s two-tailed *t*-test were used for statistical analysis. Number of experiments is indicated by *n* throughout. The data were considered as significantly different based on a confidence level of 0.05. Current measurements were plotted using ClampFit 10.2 (Molecular Devices). The IC_50_ (half maximal inhibitory concentration) and Hill coefficient (*h*) for inhibition of NMDA-evoked currents with Li^+^ were estimated by fitting of concentration-response curves with the Hill equation, I = I_min_ + (I_max_ - I_min_) /(1 + [Li^+^]^*h*^ / IC_50_^*h*^), where the I_max_ and I_min_ are the current of maximal and minimal amplitudes elicited by NMDA at different [Li+].

## Results

### Measurement of IC_50_ of lithium inhibition of NMDA-elicited currents

Extracellular Li^+^ represents a tool to cause the substrate inhibition of Na^+^-dependent Ca^2+^ extrusion by NCX. The stepwise proportional substitution of Li^+^ for Na^+^ in the bathing solution was used to obtain the dose-inhibition curve of NMDA-evoked currents for Li^+^. With this particular aim the NMDA-activated currents were measured at 0, 21, 42, 70, 112 and 140 mM Li^+^ in the bathing solution in the same experiment, where 140 mM Li^+^ corresponded to 100% substitution of Li^+^ for Na^+^. An application of Li^+^-containing solutions without agonists always preceded the application of the corresponding solution with NMDA. An increase of Li^+^ concentrations in the external solution caused a decrease of NMDA-activated current amplitude at the steady state (Fig 1A). The control NMDA-evoked currents, measured at the steady state in the bathing solutions (140 mM Na+) had the amplitude of 440.4 ± 71.9 pA (*n* = 10), that was significantly (p < 0.001, Student’s two-tailed t-test) larger compared to the corresponding value of 111.4 ± 29.1 pA (*n* = 10) measured at 140 mM Li^+^ in the external solution. Dose-inhibition curves obtained from experiments were well fitted by Hill equation with IC_50_ of 46 ± 21 mM (Fig 1B). Previously, we demonstrated that the inhibition of NMDA-activated currents by Li^+^ is Ca^2+^-dependent, because it could not be observed in the nominal absence of Ca^2+^ in the external solution [10]. Since Li^+^ does not directly affect the NMDAR conductance and activation kinetics, as a substrate inhibitor of NCXs it could sufficiently decrease the efficacy of Ca^2+^ extrusion from neurons due to breaking ion transport by NCXs. The decrease of NMDAR current amplitude by Li^+^ suggests that NCX contributes to the regulation of free Ca^2+^ concentration close to the inner membrane surface and the Ca^2+^-dependent desensitization of NMDARs. This requires some functional interaction between NCXs and NMDARs that could occur if these molecules are located closely and interact within lipid nanoclasters or rafts.

**Fig 1.**
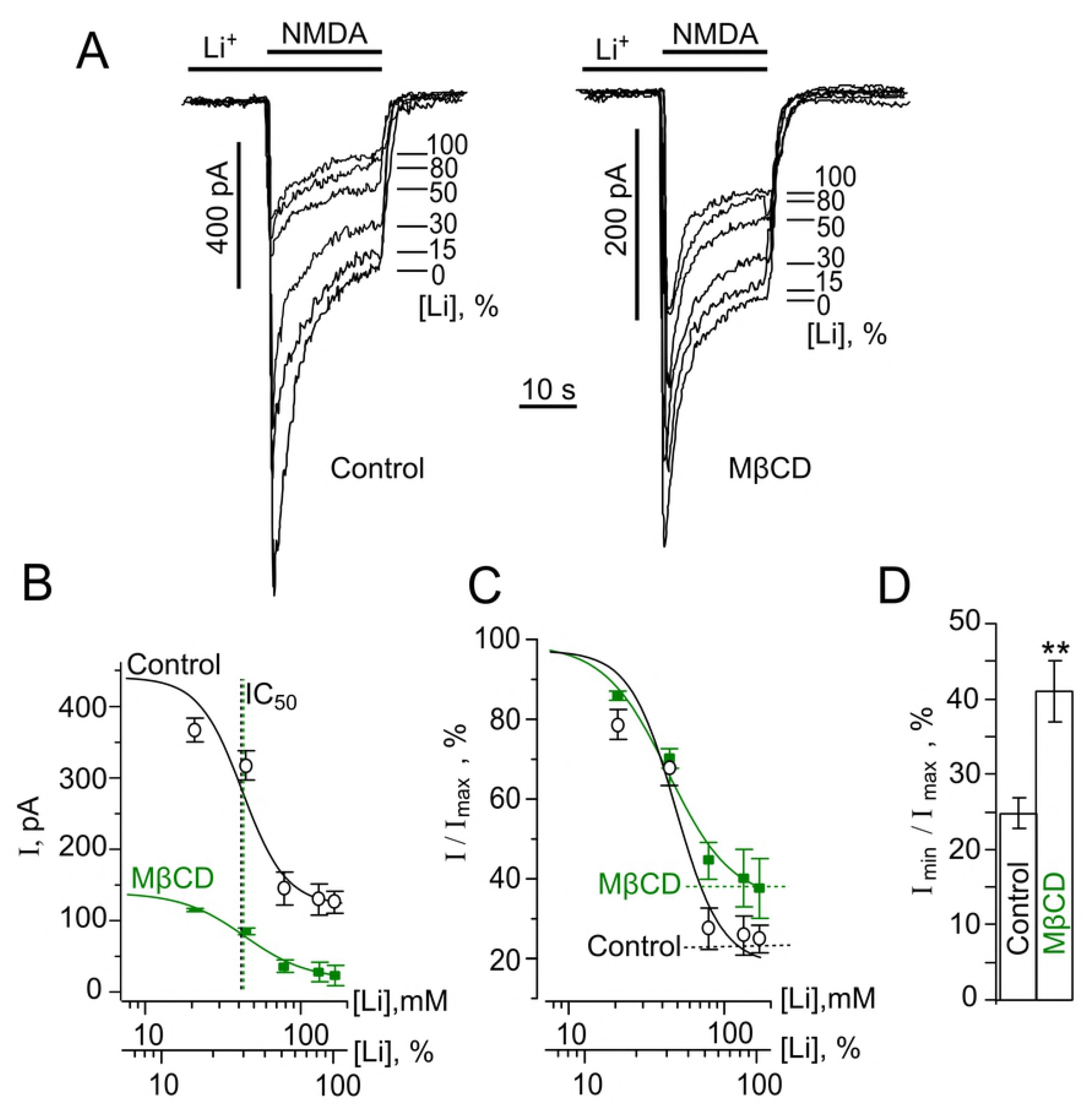
Measurements of EC_50_ of Li^+^ inhibition of NMDA-elicited currents before and after the cholesterol extraction. (A) Currents activated by 100 μM NMDA + 10 μM Gly recorded in the bathing solutions containing different Li^+^ concentrations ([Li^+^], % is indicated on the right of each trace) at – 55 mV before and after 5 min treatment with 1.5 mM MβCD. (B) Concentration-inhibition curves for Li^+^ of currents activated by 100 μM NMDA + 10 μM Gly. The mean values ± S.E.M. from 10 experiments for each of the conditions are plotted. Solid lines indicate fits of the data with the Hill equation with the parameters: IC_50_ = 46 ± 21 mM and *h* = 2.3 ± 0.8 (*n* = 10) in control conditions and 42 ± 20 mM and *h* = 3.3 ± 1.0 (*n* = 10) after the MβCD treatment. Abscissa is the Li^+^ concentration in the external solution presented as the absolute value ([Li^+^], mM) and the ratio of [Li^+^] to the sum of Na^+^ and Li^+^ concentrations of 140 mM ([Li^+^], %). (C) The same curves as on (B) normalized to I_max_ to illustrate the difference in the extent of the Li^+^ inhibition of currents, activated by NMDA before and after the MβCD treatment. (D) Histogram of fractions of residual currents (I_min_) obtained at 140 mM Li^+^ ([Li^+^], 100 %) in the external solution before (control) and after the MβCD treatment (MβCD) to the value of I_max_, drown from the fits (mean values ± S.E.M. for each of the conditions, *n* = 10). ** - the value is significantly different from the corresponding value obtained under control conditions (p < 0.01, Student’s two-tailed *t*-test).

The extraction of cholesterol from the plasma membrane by MβCD [19] is a widely used conventional procedure to destroy lipid nanoclusters. The treatment of neurons with 1.5 mM MβCD for 5 min was undertaken to extract cholesterol from membrane lipid rafts to achieve spatial uncoupling of NMDARs and NCXs. After the cholesterol extraction the mean amplitude of NMDA-evoked currents at the steady state in the Na^+^-containing bathing solution was 136.8 ± 32.8 pA (*n* = 10), revealing its decrease in comparison to MβCD untreated control conditions (p < 0.007, Student’s two-tailed *t*-test, Fig 1A and B). The stepwise substitution of Li^+^ for Na^+^ in the external solution after the MβCD treatment further decreased the NMDA-evoked currents to the mean steady-state amplitude of 46.8 ± 15.3 pA (*n* = 10, p < 0.008, Student’s two-tailed *t*-test). The IC_50_ value for the Li^+^ inhibition of NMDA-activated currents after the MβCD treatment was 42 ± 20 mM (Fig 1B and C) which did not differ significantly from the value obtained under the MβCD untreated control conditions. It should be noted, however, that the degree of the NMDAR current inhibition in the Li^+^-containing bathing solution was less pronounced after the MβCD treatment than before this procedure and were 59 ± 4 % (*n* = 10) and 77 ± 3 % (*n* = 10) inhibition (p < 0.03, Student’s two-tailed t-test), respectively (Fig 1D).

Presumably, spatial uncoupling of NCXs and NMDARs limits the effect of the NCX inhibition on NMDAR currents. This could be the case, if NCXs maintain low intracellular free Ca^2+^ concentration in the close proximity of NMDARs, which prevents the development of Ca^2+^-dependent inactivation of NMDARs.

### Calcium-dependent and –independent effects of cholesterol extraction on NMDARs

The interpretation of the above data that the cholesterol extraction may accelerate the Ca^2+^-dependent desensitization destroying membrane lipid rafts and NCX-NMDAR interplay becomes less evident in a view of the recent observation that cholesterol is important for the NMDAR functioning and its extraction provokes the ligand-dependent desensitization of NMDARs [19]. In order to distinguish between Ca^2+^-dependent and -independent mechanisms the effects of cholesterol extraction by MβCD on NMDA-activated currents were evaluated in the presence of 1 mM Ca^2+^ and in the nominal absence of Ca^2+^ in the bathing solution (Fig 2A).

**Fig 2.**
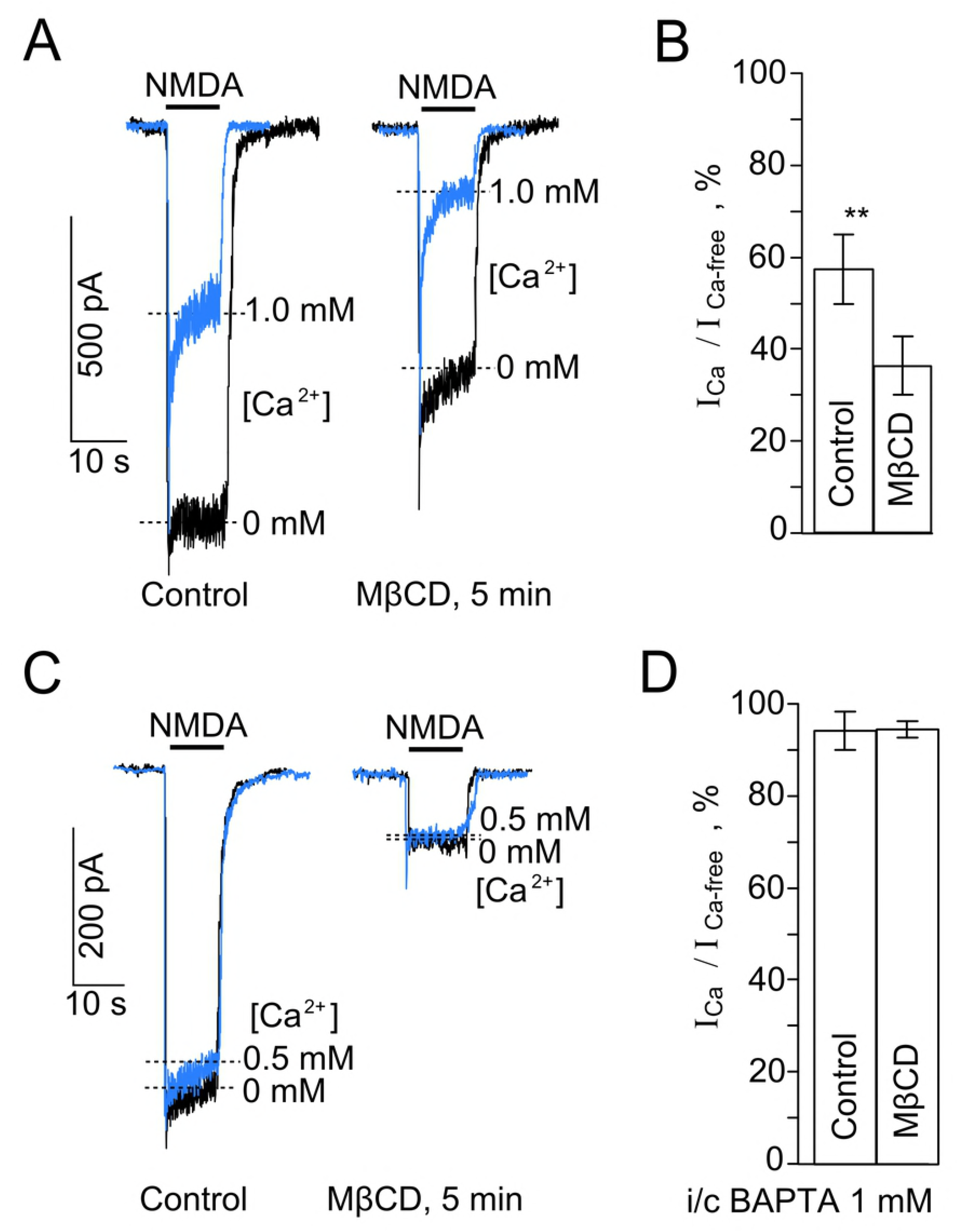
Effects of the cholesterol extraction on NMDAR currents. (A) Currents activated by 100 μM NMDA + 10 μM Gly recorded in the same neuron at – 55 mV in the nominal absence of Ca^2+^ and presence of 1 mM Ca^2+^ in the bathing solution before (Control) and after the cholesterol extraction with 1.5 mM MβCD (MβCD, 5 min). Overlays of currents are presented for the better comparison of their kinetics. (B) Comparison of NMDAR Ca^2+^-dependent desensitization before (Control) and after the cholesterol extraction (MβCD). On the histogram the amplitude ratio of currents recorded in the presence of 1 mM Ca^2+^ (I_Ca_) and the absence of Ca^2+^ (I_Ca.free_) in the external solution measured at the steady state (mean values ± S.E.M. for each of the conditions, *n* = 6). ** - the value is significantly different fro the corresponding value obtained after the cholesterol extraction (MβCD, p < 0.007, Student’s two-tailed *t*-test). (C) Currents activated by 100 μM NMDA + 10 μM Gly recorded in the same neuron at – 55 mV in the nominal absence of Ca^2+^ and presence of 0.5 mM Ca^2+^ in the bathing solution using the 1 mM BAPTA-containing intrapipette solution before (Control) and after the cholesterol extraction with 1.5 mM MβCD (MβCD, 5 min). Overlays of currents are presented for the better comparison of their kinetics. (D) Comparison of the NMDAR Ca^2+^-dependent desensitization of currents recorded using the 1 mM BAPTA-containing intrapipette solution before (Control) and after the cholesterol extraction (MβCD). On the histogram the amplitude ratio of currents recorded in the presence of 0.5 mM Ca^2+^ (I_Ca_) and the absence of Ca^2+^ (I_Ca-free_) in the external solution measured at the steady state (mean values ± S.E.M. for each of the conditions, *n* = 6).

In the absence of Ca^2+^ in the external solution the ratio of amplitudes of NMDA-activated steady-state currents, recorded after and before 5 min MβCD treatment was 47 ± 6 % (n = 6). The decrease of the NMDAR current steady-state amplitude after the treatment is caused by the direct effect of the cholesterol extraction on NMDARs, because under these particular conditions the Ca^2+^-dependent desensitization was not observed (Fig 2A). In the presence of 1 mM Ca^2+^ in the bathing solution, however, the Ca^2+^-dependent desensitization of NMDARs, measured as the ratio of the steady-state current amplitudes in the absence and presence of Ca^2+^ before and after the MβCD treatment was significantly greater when cholesterol was extracted (Fig 2A and B), suggesting that this procedure enhanced the Ca^2+^-dependent NMDAR desensitization. In addition, we performed similar experiments on neurons patched with 1 mM BAPTA in the pipette solution. Under these particular conditions the Ca^2+^-dependent desensitization of NMDARs was not observed both in the presence and absence of Ca^2+^ in the external bathing solution (Fig 2C). The direct effect of MβCD treatment on NMDARs was pronounced and the ratio values obtained in the presence and absence of external Ca^2+^ were similar (Fig 2C and D). In 1 mM intrapipette BAPTA the steady-state NMDAR currents decreased after the extraction to about 10 % of their amplitudes (Fig 2D), whereas in experiments when the intracellular media was natural in terms of Ca^2+^ buffering the NMDAR currents decreased in a much smaller extent (about 47 %, Fig 2A).

Based on these experiments we may assume that in lipid rafts NCX weakens Ca^2+^-dependent desensitization of NMDARs by quick extrusion of local intracellular Ca^2+^ entering neurons via open NMDAR pores. It is likely, that the destruction of lipid rafts increases the distance between NCXs and NMDARs allowing intracellular Ca^2+^ accumulation close to the NMDAR intracellular domains which enhances their Ca^2+^-dependent desensitization. Pronounced Ca^2+^-dependent desensitization of NMDARs, however, should provide a feed back regulation to limit the cytoplasmic Ca^2+^ accumulation during the NMDA action on neurons.

### NCX inhibition and NMDA-elicited cytoplasmic Ca^2+^ accumulation

To provide additional experimental support in favor of mechanisms suggested, online imaging of intracellular Ca^2+^ responses to 2 min NMDA applications was performed. The effects of NCX inhibition with 140 mM Li^+^ or KB-R7943 before and after the cholesterol extraction by MβCD (1.5 mM for 5 min) were studied. For quantitative comparison of effects we evaluated an integral of Ca^2+^-induced fluorescence, which has to be proportional to the Ca^2+^ entry through open NMDAR channels and, therefore, to the amplitudes of NMDA-activated currents. As in electrophysiological experiments, the Li^+^-containing bathing solution was applied alone and than with NMDA to equilibrate neurons and check pure Li^+^ effects for possible further data correction (Fig 3A). When NMDA was applied in the Li^+^-containing bathing solution Ca^2+^ responses of neurons decreased to 54 ± 2 % (n = 3, overall 98 neurons) of Ca^2+^ responses recorded in the Na^+^-containing bathing solution (p < 0.001, Student’s *t*-test). This observation is consistent with the Li^+^ effect on NMDA-activated currents. After the MβCD treatment the Ca^2+^ responses to NMDA in the Na^+^-containing solution were 35 ± 9 % (*n* = 3, overall 98 neurons) and in the Li^+^-containing solution were 36 ± 9 % (*n* = 3, overall 98 neurons) of the Ca^2+^ responses, obtained before the treatment in the Na^+^-containing solution (Fig 3A and B). Because these values were significantly smaller, than those obtained before the treatment in the Na^+^ solution (p < 0.0001, one-way paired ANOVA) and did not differ between each other (Bonferroni post-*hoc* test) we conclude that the MβCD treatment abolished the effects of Li^+^ on NMDARs.

**Fig 3.**
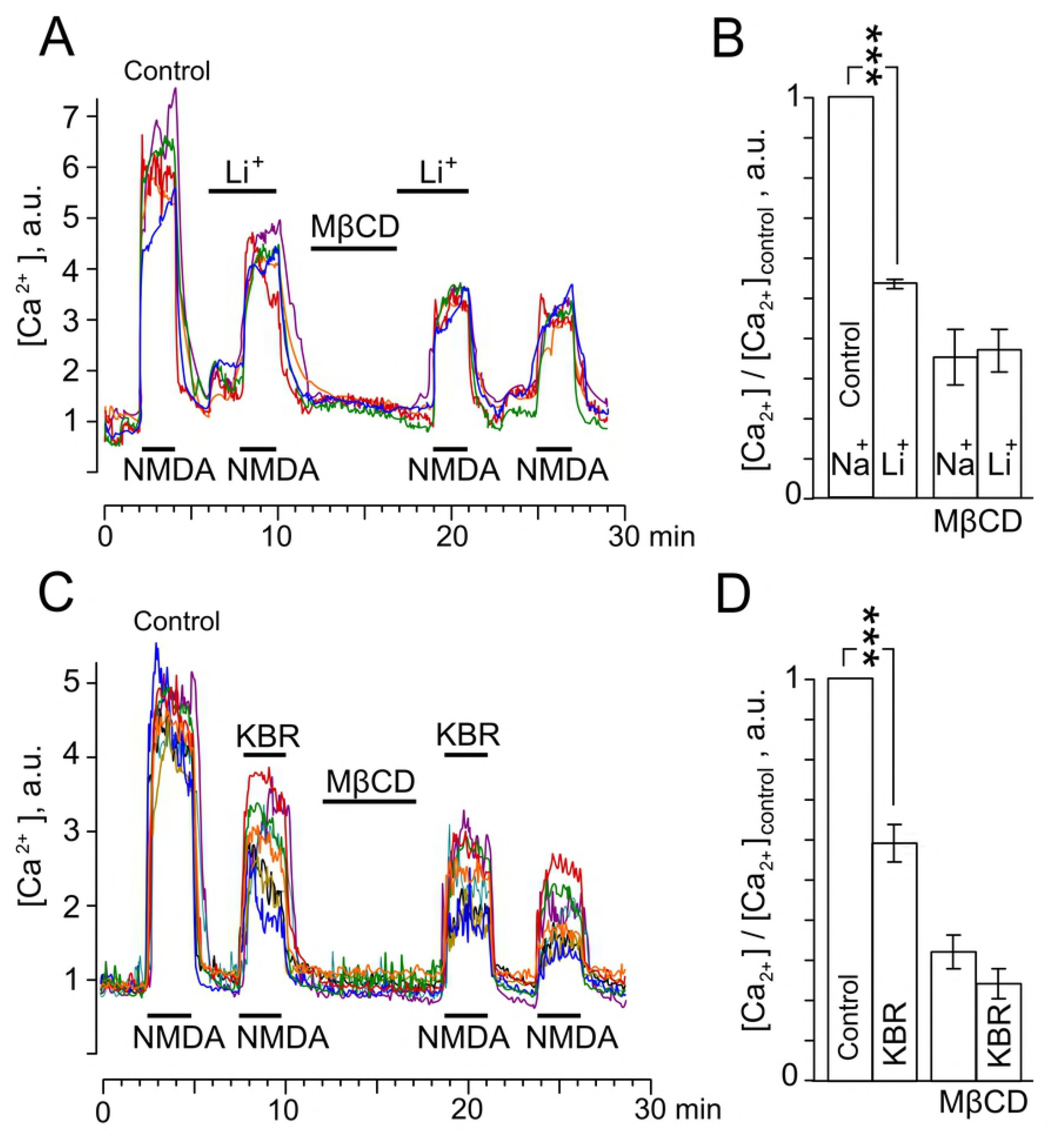
Effects of the cholesterol extraction on Ca^2+^ responses induced by NMDA. (A) Neuronal Ca^2+^ responses evoked by 100 μM NMDA ^+^ 10 μM Gly in the 140 mM Na^+^-containing (Control) and 140 mM Li^+^-containing external solutions before and after the cholesterol extraction. Applications of 100 μM NMDA + 10 μM Gly, 140 mM Li^+^ and 1.5 mM MβCD are indicated by bars. Examples of Ca^2+^ responses of 4 neurons are shown. (B) The histogram represents the ratio of squares of Ca^2+^ responses to the square of Ca^2+^ response obtained under control (140 mM Na^+^-containing external solution). Mean values ± S.E.M. for each of the conditions (*n* = 3, overall 98 neurons) are plotted. ***-the value is significantly different from other data (p < 0.0001, one-way paired ANOVA, Bonferroni post-*hoc* test). (C) Neuronal Ca^2+^ responses recorded in 140 mM Na^+^-containing external solution evoked by 100 μM NMDA + 10 μM Gly (Control) and 100 μM NMDA + 10 μM Gly + 10 μM KB-R7943 (KBR) before and after the cholesterol extraction. Applications of 100 μM NMDA + 10 μM Gly, 10 μM KB-R7943 and 1.5 mM MβCD are indicated by bars. Examples of Ca^2+^ responses of 6 neurons are shown. (D) The histogram represents the ratio of squares of Ca^2+^ responses to the square of Ca^2+^ response obtained under control (NMDA). Mean values ± S.E.M. for each of the conditions (*n* = 3, overall 91 neurons) are plotted. ***-the value is significantly different from other data (p < 0.0001, one-way paired ANOVA, Bonferroni post-*hoc* test).

Thus, spatial uncoupling of NMDARs and NCXs resulted in the decrease of Ca^2+^ entry via NMDARs. Inhibition of NCX with Li^+^ after the cholesterol extraction was not able to decrease NMDARs mediated Ca^2+^ accumulation.

We further performed the Ca^2+^ imaging experiments in which KB-R7943 (10 μM) as a specific NCX inhibitor, was utilized instead of the Li^+^ solution. In the Na^+^-containing external solution combined applications of NMDA with KB-R7943 induced Ca^2+^ responses that corresponded to 59 ± 5 % (n = 3, overall 91 neurons) of NMDA-elicited Ca^2+^ responses and differed from them significantly (p<0.001, one-way paired ANOVA) (Fig 3C). This observation is consistent with the KB-R7943 effect on NMDA-activated currents [10]. The MβCD treatment decreased the NMDA-elicited Ca^2+^ responses both in the absence and presence of KB-R7943 to 32 ± 8 % and 24 ± 7 % (*n* = 3, overall 91 neurons), respectively (Fig 3D). These values are not significantly different (Bonferroni post-*hoc* test) suggesting that the cholesterol extraction abolished the KB-R7943 effects on NMDA-activated currents.

Therefore, the effects of Li^+^ and KB-R7943 on NMDA-elicited Ca^2+^ responses of neurons coincide well suggesting that they both are realizing through the influence of NCX on the Ca^2+^-dependent desensitization of NMDARs.

## Discussion

In spite of a large number of novel pharmacological agents has recently appeared, Li^+^ has still a broad usage as a tool of neuroscience researches, since it can affect key functional processes of the central nervous system (CNS) including different enzymes [for review, see 12] and Na^+^-dependent neurotransmitter transporters [11] and exchangers [for review, see 13]. Diverse and complex action of Li^+^ on the human CNS is highlighted by the Li^+^ therapy which is widely utilized to stabilize many mental disorders. Usually therapeutic Li^+^ concentrations in the blood vary within the range of 0.6 - 1.2 mM and the concentrations over 1.5 mM are thought to become toxic [for review, see 12]. In addition it has been demonstrated recently that the substitution of Li^+^ for Na^+^ in the external solution in the presence of Ca^2+^ causes considerable decrease of currents activated by NMDA [10]. This somehow contradicts to the lack of the NMDAR Li^+^ inhibition in the absence of Ca^2+^ in the external solution [10] and to the observation that Li^+^ does not influence the NMDAR conductance and activation kinetics [14]. The IC_50_ value for the Li^+^ inhibition of NMDA-activated currents measured here is about 44 mM. It is, therefore, unlikely that the Li^+^ inhibition of NMDAR currents in some extent contributes in the therapeutic effect during the Li^+^ therapy, whereas the mechanism of the Ca^2+^-dependent Li^+^ inhibition of NMDARs requires further consideration.

The critical dependence of Li^+^ inhibition of NMDAR currents on extracellular Ca^2+^ forced us to the conclusion that Li^+^ inhibits NMDARs indirectly breaking the Ca^2+^ extrusion from neurons by NCXs, which are involved in regulation of pre-membrane Ca^2+^ concentration in the close proximity to the NMDAR intracellular domains during Ca^2+^ entry through the channels of activated NMDARs. By the other words Li^+^ promotes Ca^2+^-dependent desensitization of NMDARs inhibiting the NCX transport of Ca^2+^ from neurons [10]. If this is the case then NMDARs and NCXs should co-localize and interact that could be achieved in membrane cholesterol rich nanocluster or lipid rafts (Fig. 4A). Actually the co-localization of NMDARs and NCXs in lipid rafts at the distance less than 80 nm was recently demonstrated using FRET (Förster Resonance Energy Transfer) experiments [20, 21]. In our experiments the cholesterol extraction, that is known to destruct lipid rafts, resulted in a substantial decrease of NMDAR currents, which is consistent to the earlier observation [19], but did not cause significantly changes of the IC_50_ value for the Li^+^ inhibition of NMDAR currents. This may suggest that the cholesterol extraction does not influence the transport by NCXs [22] and similar Li^+^ concentrations are required to inhibit CNXs before and after the MbCD treatment.

**Fig 4.**
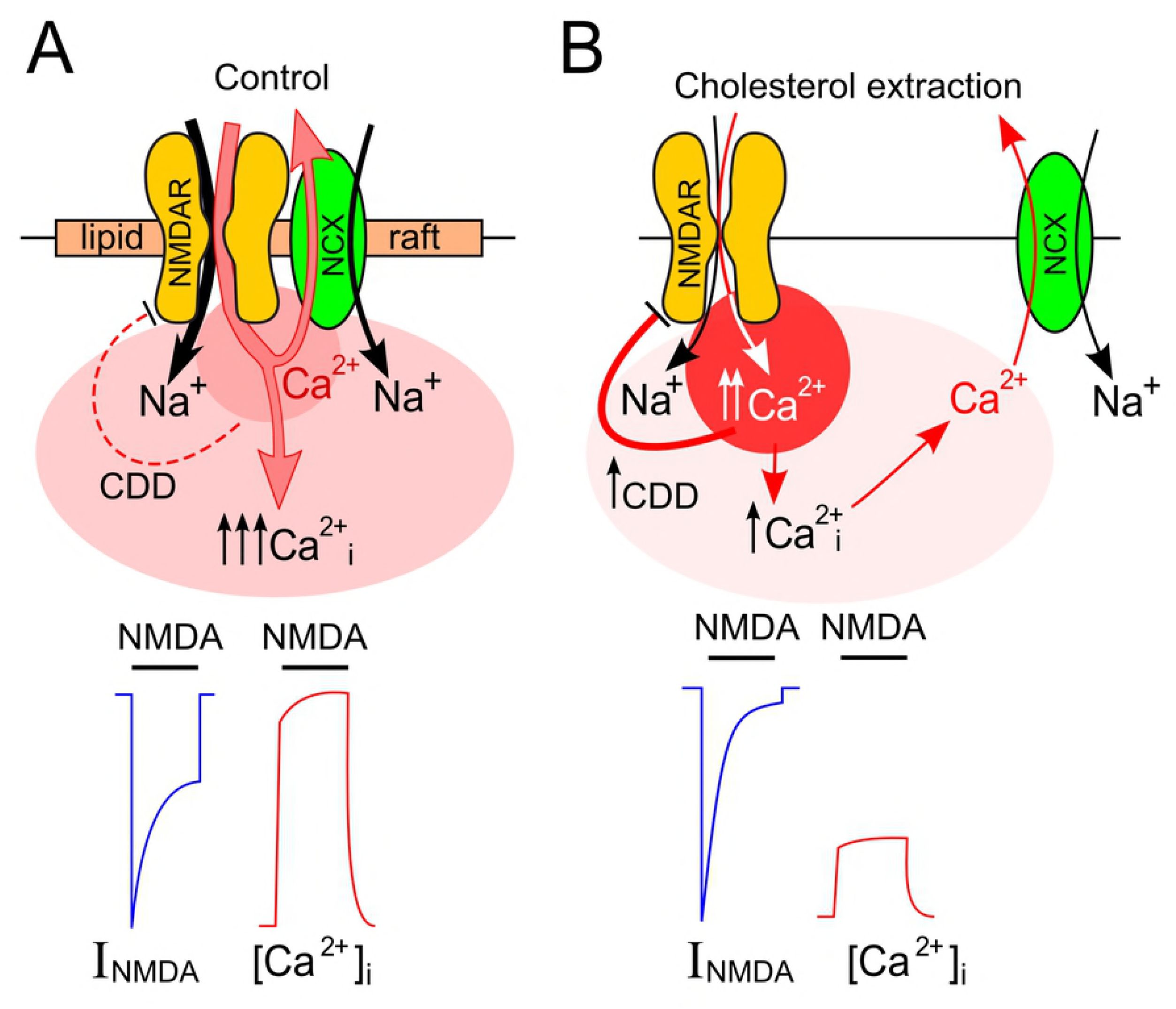
Schematics of the data interpretation. (A) In control conditions local Ca^2+^ accumulation in the close proximity to intracellular domains of activated NMDARs is prevented by NCX-transported Ca^2+^ extrusion. The integral Ca^2+^ entry into neurons via activated NMDARs is large because of moderate Ca^2+^-dependent NMDAR desensitization (CDD). This causes prominent cytozolic [Ca^2^+]_i_ increase (↑↑↑Ca^2+^_i_). Typical whole-cell current (I_NMDA_) and fluorescent Ca^2+^ probe response are indicated below the panel. (B) The destruction of lipid rafts increases a distance between NMDARs and NCXs which prevents fast removal of Ca^2+^ entering via open NMDAR channels. Local Ca^2+^ accumulation in the close proximity of intracellular domains of activated NMDAR enhances their CDD and limits NMDAR current steady-state amplitude and as a consequence weakens total cytosolic Ca^2+^ accumulation. Typical whole-cell current (I_NMDA_) and fluorescent Ca^2+^ probe response ([Ca^2+^]_i_) are indicated below the panel. For further explanation, see Discussion.

The requirement of cholesterol for functioning of NMDARs was recently demonstrated [19], because its extraction induced fast ligand-dependent NMDAR desensitization. In agreement when we used 1 mM BAPTA containing intrapipette solution (calculated free Ca^2+^ concentration is 13 nM) a 10-fold decrease of NMDAR currents and a lack of Ca^2+^-dependent NMDAR desensitization were observed. NMDA-activated currents recorded in the absence of Ca^2+^ in the external solution using the BAPTA-free intrapipette solution did not reveal the NMDAR Ca^2+^-dependent desensitization as well. The cholesterol extraction under these conditions, however, induced a 2-fold decrease of the NMDAR currents. The lesser extent of the ligand-dependent desensitization obtained without BAPTA may suggest that some normal level of free Ca^2+^ in the cytoplasm is required for NMDAR functioning. In addition the extraction caused an enforcement of Ca^2+^-dependent NMDAR desensitization suggesting that the disaggregation of molecules within destructed lipid rafts is accompanied by the disruption of NCX regulated Ca^2+^ pre-membrane balance.

Measurements of intracellular Ca^2+^ dynamics revealed that the NCX inhibition with Li^+^ or KB-R7943 significantly decreased the NMDA-elicited Ca^2+^ responses, which is consistent to their effects on currents, activated by NMDA. In agreement to previous observations [23] the cholesterol extraction caused the decrease of cytoplasm Ca^2+^ accumulation, and furthermore abolished the effects of both Li^+^ and KB-R7943 on neuronal Ca^2+^ cytoplasmic responses. These further support our conclusion that the destruction of lipid rafts abolishes the influence of NCXs on NMDARs (Fig. 4B).

These experiments allow us to suggest, that the NCX inhibition prevents the maintainance of low Ca^2+^ level in the proximity of the intracellular domains of NMDARs by the Ca^2+^ extrusion to the outside, which elevates pre-membrane local Ca^2+^ concentration, but limits total Ca^2+^ entry into neurons. Modeling of interaction between CaM and C-terminal of GluN1 subunits of NMDARs reveals that it could occur if these molecules are co-localized within the distance of tens of nanometers [24]. Spatial uncoupling of NCXs and NMDARs weakens the NCX influence on the Ca^2+^-dependent NMDAR desensitization. Thus, the inhibition of NCXs with Li^+^ or KB-R7943 after the disaggregation of molecules within former lipid rafts does not significantly influence the cytoplasmic Ca^2+^ accumulation in response to NMDAR activation.

Author Contributions
Conceptualization: SMA, DAS
Experimental work: DAS, EEP
Data analysis and plotting: DAS, EEP, SMA
Writing the manuscript: DAS, SMA
Funding acquisition: SMA

